# Timing and location of speech errors induced by direct cortical stimulation

**DOI:** 10.1101/2023.09.14.557732

**Authors:** Heather Kabakoff, Leyao Yu, Daniel Friedman, Patricia Dugan, Werner K Doyle, Orrin Devinsky, Adeen Flinker

## Abstract

Cortical regions supporting speech production are commonly established using neuroimaging techniques in both research and clinical settings. However, for neurosurgical purposes, structural function is routinely mapped peri-operatively using direct electrocortical stimulation. While this method is the gold standard for identification of eloquent cortical regions to preserve in neurosurgical patients, there is lack of specificity of the actual underlying cognitive processes being interrupted. To address this, we propose mapping the temporal dynamics of speech arrest across peri-sylvian cortices by quantifying the latency between stimulation and speech deficits. In doing so, we are able to substantiate hypotheses about distinct region-specific functional roles (e.g., planning versus motor execution). In this retrospective observational study, we analyzed 20 patients (12 female; age range 14-43) with refractory epilepsy who underwent continuous extra-operative intracranial EEG monitoring of an automatic speech task during clinical bedside language mapping. Latency to speech arrest was calculated as time from stimulation onset to speech arrest onset, controlling for individual speech rate.

Most instances of motor-based arrest (87.5% of 96 instances) were in sensorimotor cortex with mid-range latencies to speech arrest with a distributional peak at 0.47 seconds. Speech arrest occurred in numerous regions, with relatively short latencies in supramarginal gyrus (0.46 seconds), superior temporal gyrus (0.51 seconds), and middle temporal gyrus (0.54 seconds), followed by relatively long latencies in sensorimotor cortex (0.72 seconds) and especially long latencies in inferior frontal gyrus (0.95 seconds). Nonparametric testing for speech arrest revealed that region predicted latency; latencies in supramarginal gyrus and in superior temporal gyrus were shorter than in sensorimotor cortex and in inferior frontal gyrus. Sensorimotor cortex is primarily responsible for motor-based arrest. Latencies to speech arrest in supramarginal gyrus and superior temporal gyrus (and to a lesser extent middle temporal gyrus) align with latencies to motor-based arrest in sensorimotor cortex. This pattern of relatively quick cessation of speech suggests that stimulating these regions interferes with the outgoing motor execution. In contrast, the latencies to speech arrest in inferior frontal gyrus and in ventral regions of sensorimotor cortex were significantly longer than those in temporoparietal regions. Longer latencies in the more frontal areas (including inferior frontal gyrus and ventral areas of precentral gyrus and postcentral gyrus) suggest that stimulating these areas interrupts a higher-level speech production process involved in planning. These results implicate the ventral specialization of sensorimotor cortex (including both precentral and postcentral gyri) for speech planning above and beyond motor execution.

## Introduction

Direct electrocortical stimulation (DES) mapping is routinely used peri-operatively in patients with epilepsy and other brain anomalies to functionally map regions critical for motor, sensory, speech, and language functions.^1^ Cortical stimulation results allow clinicians to map resection boundaries and avoid post-operative deficits, representing the gold standard to prevent functional impairments following epilepsy surgery.^2,3^

During DES mapping, cortex is stimulated while patients perform speech tasks. At our institution, these tasks include recitation of automatic speech, visual/auditory naming, and sentence completion,^1,4^ though tasks such as those eliciting continuous speech and visual naming are shared by many institutions. Hesitation, slurring, distortion, repetition, and _1,2_ confusion can interrupt any task. We distinguish “motor-based arrest” from “speech arrest” based on an established clinical criterion. Both motor-based arrest and speech arrest occur when there is complete cessation of speech during a continuous speaking task. We define motor-based arrest as the kind of speech cessation that occurs when stimulation impairs control within the vocal tract, as evidenced by unintended oral, pharyngeal, or laryngeal movements; therefore, we define speech arrest as the complete interruption of speech that cannot be explained by motor interruption.^5,6^

Based on classical^1^ and recent reports^6,7^ on speech arrest sites, the precentral gyrus is the area most commonly reported to induce speech arrest during DES, followed by the inferior frontal gyrus (IFG).^1,6-8^ Recent statistical probability distributional maps converge on precentral gyrus and IFG pars opercularis as the two regions with the highest statistical probability of inducing speech arrest.^7,8^ Of particular relevance to this study, functional maps derived from DES have indicated that electrodes in precentral gyrus and postcentral gyrus produce comparable speech arrest patterns, particularly in ventral portions.^6,9^

In these same intra-operative studies, the superior temporal gyrus (STG) is less commonly and less consistently reported as being associated with speech arrest than precentral gyrus or IFG.^1,6,7^ Statistical probability maps have identified STG and the supramarginal gyrus to form the fourth largest cluster with the highest probability of inducing speech arrest, after the largest clusters in precentral gyrus and pars opercularis.^6^ STG inconsistently induced speech arrest across patients,^8^ with great variability between subjects,^7^ suggesting that STG is strongly associated with speech arrest in some speakers and not others.

Other inconsistencies occur in cortical regions in which stimulation causes speech arrest. Two studies in particular found robust speech arrest in precentral gyrus.^5,9^ However, one of these studies found sparse speech arrest in IFG (4% chance of speech arrest), STG (0%), and supramarginal gyrus (8%).^9^ The other study found sparse speech arrest in IFG or STG, and none in supramarginal gyrus.^5^ Additionally, there have been reports of regions inducing speech arrest even though they were not classically reported. First, statistical probability maps identified an isolated area in the supplemental motor area as the third largest cluster with the highest probability of inducing speech arrest.^6^ Finally, two DES studies have reported speech arrest in ventral portions of postcentral gyrus^6,9^ even though sensorimotor electrodes are generally limited to precentral gyrus. Some of these inconsistencies likely reflect clinical differences in mapping procedures (e.g., differences between intra-operative DES^1,6-9^ versus extra-operative DES mapping^5^) and differences in coverage for epilepsy^1,5,7,8^ versus tumor mappings.^6,8,9^ Differences in stimulation parameters, particularly the electrical current intensity, within and across sites may also contribute to inconsistent prior findings.

As in past studies, we also report the spatial location of speech arrest. However, the goal of the present study is to explore timing and make associated conclusions about the functional roles of regions. We do this by measuring the timing of how long it takes stimulation to induce speech arrest in five peri-sylvian cortical regions during extra-operative DES mapping. A previously reported method for measuring timing relative to speech interruptions involved using EEG data to calculate “post-ictal language delays,” or the time (in seconds) from the offset of spontaneous seizure activity until a patient continued speaking.^10^ While this characterizes the temporal relationship between abnormal (i.e., epileptic) activity and speech interruptions, it does not inform the causal connection between cortical sites and speech production, for which the DES method is specifically designed. Further, intracranial electrodes provide maximized spatial specificity compared to EEG. Here, we leverage clinically-induced interruptions following cortical stimulation (while ruling out epileptic activity) as our approach for measuring timing.

One prior study employing repetitive transcranial magnetic stimulation (rTMS) reported that the timing from stimulation onset to speech arrest in left precentral gyrus was 1±3s.^11^ A related prior study applied transcranial magnetic stimulation (TMS) at various time points following stimulus presentation in a picture naming task.^12^ They identified that stimulation at 225 ms over MTG, at 300 ms over IFG, and at 400 ms over STG led to longer latencies to production than stimulation in the same regions at other time points. This suggests that stimulation interrupted distinct stages of speech production in these three regions.^12^ While this past study reported the timing of speech interruption following TMS, we are unaware of any past studies reporting the timing of speech arrest following DES in order provide a window into the functional architecture and neural timing underlying speech production. Using DES during extra-operative clinical mapping, our data set offers unprecedented spatial and temporal precision to distinguish latencies in different cortical regions. Mapping out latencies for inducing speech arrest across cortex has strong potential to inform which phase in the speech production process various types of interruptions occurred.

We assess whether latencies from stimulation onset to speech arrest during a continuous speaking task differ across five peri-sylvian cortical regions: sensorimotor cortex (which we define anatomically to include both precentral and postcentral gyri, following the example set in recent research^13-15^), IFG, STG, supramarginal gyrus, and middle temporal gyrus (MTG). Speech production models highlight motor cortex in executing motor programs and IFG in linking phonology to speech motor control.^16,17^ During word repetition, neural representations are forwarded from STG to IFG, where they link to articulatory representations, which are subsequently implemented by the articulators via motor cortex. Since IFG activity precedes speech production^18-20^ by approximately 250 milliseconds,^18^ we hypothesized that stimulation in sensorimotor cortex would more rapidly interrupt speech than IFG stimulation. We predicted that sensorimotor cortex stimulation would interrupt mid-trajectory in the motor execution phase of speech production, resulting in the shortest latencies to speech arrest.

Conversely, we predicted that IFG stimulation would cause interruptions to the outgoing plan, which should only affect future speech, resulting in relatively long latencies from stimulation to speech arrest. These hypotheses are consistent with sensorimotor cortex and inferior frontal gyrus (IFG) being the most likely cortical sites to elicit speech arrest, as well as their putative functional roles in speech production models.^16,18,21,22^ Therefore, we would interpret relatively short latencies to speech arrest following stimulation in sensorimotor cortex and relatively long latencies in IFG as corroborating hypotheses about speech production models that separate planning and execution processes anatomically.

## Materials and methods

### Subject selection

All the patients at NYU Langone Hospital who underwent DES mapping and had precise synchronization of EEG with audio in the clinical system were considered for inclusion in this retrospective observational study. This comprised fifty patients who were implanted from September 2018 through September 2021, beginning from when data were available in the Natus® Neuroworks® EEG system, providing a software development kit (SDK) to synchronize audio and EEG with high fidelity. All patients were implanted with subdural and depth electrodes (AdTech Medical Instrument Corp.) and underwent bedside extra-operative DES mapping to identify eloquent (motor, language) cortex prior to surgical resection of epileptogenic tissue as part of their routine clinical care. A total of 20 patients (12 females; mean age 27.7, SD = 10.46, age range = 14-43; 1 right, 19 left hemisphere coverage; mean age of diagnosis 12.4, SD = 8.0) were reported to have speech interruptions that were labeled as motor-based or as language-based (including “speech arrest”). This determination will be further defined in a later section. Electrode coverage typically comprised grids (8×8 contact, 10 mm inter-electrode distances), strips over cortex (4-8 electrodes, 10 mm inter-electrode distance), and depth electrodes (≤8 electrodes, 5 mm inter-electrode distance). Surface reconstructions, electrode localization, and Montreal Neurological Institute (MNI) coordinates were extracted by aligning a postoperative brain MRI to the preoperative brain MRI using previously published methods.^23^ See plots of electrode coverage for all patients in Supplemental Fig. S1.

### Informed consent

All patients gave informed consent to participate in research and for audio/video to be recorded and analyzed, as approved by the New York University Langone Health Institutional Review Board. Table 1 lists all twenty patients considered for this analysis.

**Table 1.**
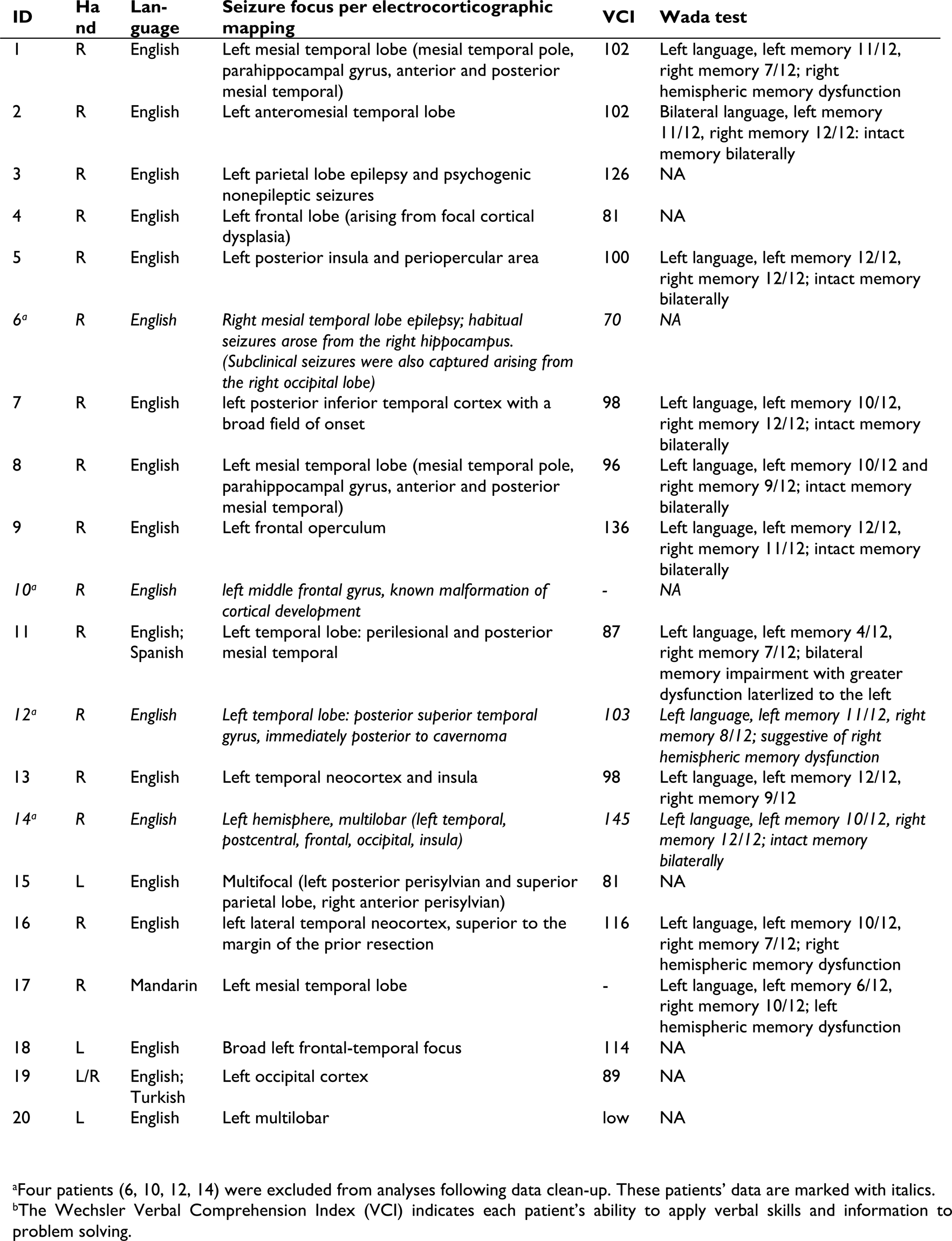
Patient characteristics, including anonymized ID, handedness, languages spoken ordered by proficiency, seizure focus, verbal skill, and language/memory lateralization for all twenty originally included patients.

| ID | Hand | Language | Seizure focus per electrocorticographic mapping | VCI | Wada test |
| --- | --- | --- | --- | --- | --- |
| 1 | R | English | Left mesial temporal lobe (mesial temporal pole, parahippocampal gyrus, anterior and posterior mesial temporal) | 102 | Left language, left memory 11/12, right memory 7/12; right hemispheric memory dysfunction |
| 2 | R | English | Left anteromesial temporal lobe | 102 | Bilateral language, left memory 11/12, right memory 12/12; intact memory bilaterally |
| 3 | R | English | Left parietal lobe epilepsy and psychogenic nonepileptic seizures | 126 | NA |
| 4 | R | English | Left frontal lobe (arising from focal cortical dysplasia) | 81 | NA |
| 5 | R | English | Left posterior insula and periopercular area | 100 | Left language, left memory 12/12, right memory 12/12; intact memory bilaterally |
| 6<sup>a</sup> | R | English | *Right mesial temporal lobe epilepsy; habitual seizures arose from the right hippocampus. (Subclinical seizures were also captured arising from the right occipital lobe)* | 70 | NA |
| 7 | R | English | left posterior inferior temporal cortex with a broad field of onset | 98 | Left language, left memory 10/12, right memory 12/12; intact memory bilaterally |
| 8 | R | English | Left mesial temporal lobe (mesial temporal pole, parahippocampal gyrus, anterior and posterior mesial temporal) | 96 | Left language, left memory 10/12 and right memory 9/12; intact memory bilaterally |
| 9 | R | English | Left frontal operculum | 136 | Left language, left memory 12/12, right memory 11/12; intact memory bilaterally |
| 10<sup>a</sup> | R | English | *left middle frontal gyrus, known malformation of cortical development* | - | NA |
| 11 | R | English; Spanish | Left temporal lobe: perilesional and posterior mesial temporal | 87 | Left language, left memory 4/12, right memory 7/12; bilateral memory impairment with greater dysfunction lateralized to the left |
| 12<sup>a</sup> | R | English | *Left temporal lobe: posterior superior temporal gyrus, immediately posterior to cavernoma* | 103 | *Left language, left memory 11/12, right memory 8/12; suggestive of right hemispheric memory dysfunction* |
| 13 | R | English | Left temporal neocortex and insula | 98 | Left language, left memory 12/12, right memory 9/12 |
| 14<sup>a</sup> | R | English | *Left hemisphere, multilobar (left temporal, postcentral, frontal, occipital, insula)* | 145 | *Left language, left memory 10/12, right memory 12/12; intact memory bilaterally* |
| 15 | L | English | Multifocal (left posterior perisylvian and superior parietal lobe, right anterior perisylvian) | 81 | NA |
| 16 | R | English | left lateral temporal neocortex, superior to the margin of the prior resection | 116 | Left language, left memory 10/12, right memory 7/12; right hemispheric memory dysfunction |
| 17 | R | Mandarin | Left mesial temporal lobe | - | Left language, left memory 6/12, right memory 10/12; left hemispheric memory dysfunction |
| 18 | L | English | Broad left frontal-temporal focus | 114 | NA |
| 19 | L/R | English; Turkish | Left occipital cortex | 89 | NA |
| 20 | L | English | Left multilobar | low | NA |

<sup>a</sup>Four patients (6, 10, 12, 14) were excluded from analyses following data clean-up. These patients' data are marked with italics.
<sup>b</sup>The Wechsler Verbal Comprehension Index (VCI) indicates each patient's ability to apply verbal skills and information to problem solving.

### Clinical battery

As part of the battery of clinical tasks routinely administered to elicit language,^24^ the current focus is on the continuous speech elicited during counting or reciting days of the week, months of the year, or the Pledge of Allegiance. Though not analyzed in the present study, patients also completed higher-level tasks eliciting visual and auditory naming and auditory comprehension with the goal of capturing multiple modalities of language processing, as described in our previous research.^25^ Following clinical protocol,^24^ stimulation was presented to electrode pairs using a NicoletOne Cortical Stimulator which typically involved bipolar contiguous contacts delivering a low-intensity current (0.5-6 mA) that was gradually increased (maximum 15 mA) across multiple trials to find the threshold at which cortical spread (afterdischarges) did not occur and a behavioral deficit was observed. Current was delivered with a biphasic pulse width of 300-500 μs at a pulse rate of 50 Hz with a train duration from 3-5 s. An epileptologist inspected each channel and trials eliciting afterdischarges were labeled as such for future consideration (see below).

Following clinical protocol at NYU Langone, for each electrode pair (dipole 1 and dipole 2), each patient began counting as their voice was recorded using an external microphone connected to the clinical audio/video system. Stimulation was delivered in selected intervals during continuous automatic speech, beginning at the lowest intensity current level and gradually increasing within established thresholds until a behavioral deficit was observed. Although there were multiple stimulation trials within a sequence that were delivered at different stimulation intensity levels, only the level for the final trial was documented following clinical protocol (see results and discussion). Once a behavioral deficit was observed, stimulation was repeated while the patient continued the speaking task to see whether the same deficit occurred on at least two out of three trials. When there was no behavioral deficit, the target electrode pair was considered “cleared” as not controlling movement or language. If the site was found to be a hit for having a motor-based cause (i.e., “motor hit,” see next paragraph for how this was determined), then stimulation mapping was complete for that electrode pair. If the site was determined to be a hit for having a language-based cause (“language hit”), then a higher-level continuous speaking task was elicited (e.g., days of the week, months of the year, Pledge of Allegiance) to probe further for speech interruptions. We acknowledge that the first three tasks (counting, reciting days/months) can be considered more automatic than the Pledge of Allegiance, which would be considered to involve more propositional language. However, this task was only used to elicit speech from those who had it memorized, suggesting that it was produced with a high degree of automaticity.

Based on the epileptologist reports available in the clinical system, individual electrodes that elicited a behavioral response were labeled as “motor hits” or as “language hits.” This determination was made by first identifying that speech was interrupted and then ruling out whether each speech interruption had a motor-based cause. Distinct from the continuous speaking task in which automatic speech was elicited (counting, reciting days/months/Pledge), whether a speech interruption had a motor-based cause was confirmed separately using a nonspeech task involving rapid repeated production of syllables (i.e., diadochokinesis). A stimulation site was determined to be a motor hit if a part of the patient’s body reliably moved during diadochokinesis, thus interrupting speech output. Specifically, a patient’s overt lip or tongue movement interfering with sequential production “pa,” “ta,” “la”, or “ka” would be considered a motor-based speech interruption (“motor hit”). As laryngeal movement is more covert than oral movement, patient report of feeling movements stuck in their throat would be interpreted by the clinician as motor hits with laryngeal involvement. Interruptions were only considered motor hits if the motor activity directly interfered with speech output. Therefore, through the process of elimination, those speech interruptions that were not motor hits were determined to be caused by interruption to higher-level language processes, and were therefore called “language hits.” We define “speech arrest” as the complete interruption of the ability to continue speaking that is not directly explained by oral, pharyngeal, or laryngeal movements.^5,6^ As such, only those language hits involving complete cessation of speech were subcategorized as speech arrest.

### Measurement

Using Neuroworks, we viewed the video recordings of DES bedside mapping for all 20 patients. Time stamps for the onset of stimulation for all speech interruptions during continuous speaking were noted in conjunction with dipole electrode labels. Although our focus is on speech interruptions independent of whether they are due to motor causes, we liberally included in the initial data set all trials involving a noticeable change in speech output coinciding with stimulation.

To obtain precise recordings of stimulation intervals, we read in proprietary Natus files (.ERD, .ETC., .STC, .SNC) and extracted the EEG signal from the first in each pair of target electrodes (dipole 1) using custom Matlab^26^ code based on the SDK provided by Natus. For each patient, we read in the Natus video file and extracted the time-synced audio signal from the bedside recording (synchronization was based on the associated synchronization files used by Natus, including .ETC, .STC, .SNC, and .VTC formats). For each patient and unique target electrode (dipole 1), we extracted a stereo audio recording including stimulation signals (channel 1) and time-synced audio (channel 2).

Using Praat software^27^ for acoustic analysis, we generated time-synced TextGrids for each extracted stereo audio file. Fig. 1 (panel A) shows an example audio signal and aligned annotated TextGrid, including one speech arrest event. First, a window was created around each speech interruption, which encompassed utterances leading up to and following the interruption up until the next stimulation interval began or the clinician intervened. Within those windows, a tier (“word”) was used to label the onset and offset of each word in each target sequence. On a second tier (“stim”), we annotated the precise timing of stimulation, which was visible as each interval of non-zero amplitude in the EEG signal from channel 1. A final tier (“arrest”) was used to mark the moment at which speech interruptions occurred, including complete cessation of speech (marked with an asterisk). Also coded on the “arrest” tier, symbols denote whether the patient was cut off from continuing speaking (+/-) and whether the patient could continue where they left off (c) following the speech interruption. Though not reported presently, additional observed changes in the speech output were noted on this tier, including slowed speech, pauses, non-target vocalizations, perseverations or skips in the sequence, changes in vocal quality or volume, and speech aberrations classified as dysarthric or apraxic.

**Figure 1.**
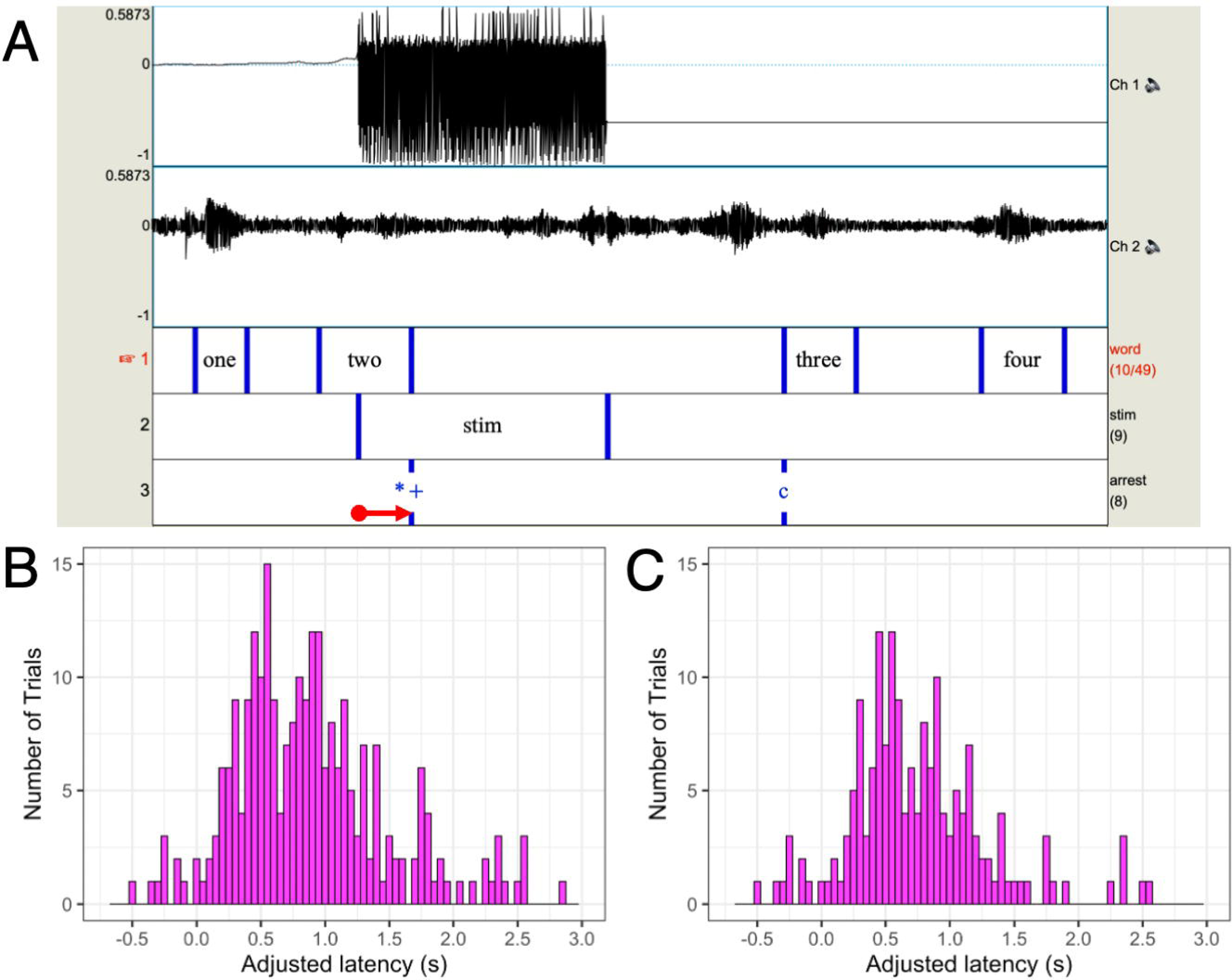
Measurement and distributions of adjusted time elapsed since stimulation onset, controlling for trials with afterdischarges. **(A)** In Praat acoustic analysis software, we visualized the stimulation intervals (channel 1) and the audio signal (channel 2). Based on channel 1, we annotated precise onset and offset of stimulation in a tier labeled “stim.” Within each trial in channel 2, we annotated each word interval in a tier labeled “word,” and the exact timing of speech arrest events in a tier labeled “arrest.” Symbols on the “arrest” tier denote speech arrest (*), that the patient was able to continue speaking after the initial interruption (+), and that they continued counting right where they left off (c). Raw latencies, as shown with the red arrow, were calculated as the time (in seconds) elapsed between stimulation onset (from the “stim” tier) and the onset of speech arrest (from the “arrest” tier). Adjusted latencies were derived by adding to the raw latencies the average duration between all words prior to stimulation onset in each trial sequence. **(B)** Distribution of all adjusted latencies, including trials with afterdischarges. **(C)** Distribution of all adjusted latencies, excluding trials with afterdischarges.

The primary outcome measure is latency from stimulation onset to speech arrest. One limit of analyzing speech arrest timing is ambiguity of interruption onset, even in continuous speech. In contrast to speech degradations that can be marked in the audio, speech arrest onset corresponds with signal loss. If this does not occur suddenly in the middle of a word, it may occur at some point between the end of the last word spoken and the time when the next word would have been spoken at the same rate. As true latencies cannot be directly measured because of these pauses, we used the following method to approximate them. Raw latencies were calculated as the time from the onset of stimulation to the onset of speech arrest (see Fig. 1, panel A). Thus, negative latencies are possible because speech arrest is marked at the end of the last word that was spoken, which could occur before stimulation onset even though the true speech arrest would be prior to the next expected word in the sequence. To account for pre-stimulation speech rate within and between patients, we adjusted raw latencies by adding the average duration between all words prior to stimulation onset in each trial (and between phrases for the Pledge of Allegiance). Making these adjustments to the raw latencies eliminated many negative latencies while providing the closest possible latency estimations.

Timings were extracted from the measured Praat TextGrid boundaries and read into R^28^ using RStudio.^29^ Each electrode label was matched with both the name of the corresponding brain region based on the subject’s own pre-operative MRI and the standardized MNI coordinates. Each speech interruption was matched with the label “motor hit” or “language hit” based on the clinician reports. As previously described, electrode pairs determined to be language hits were also considered speech arrest hits if the clinical report listed that electrode pair as a hit for speech arrest. As multiple trials were elicited for each electrode pair, only those trials with a hard cessation of speech were labeled as speech arrest.

### Analysis

Across the 20 patients, there were 360 trials labeled as motor hits or speech arrest hits by the clinical team. Latencies greater than two standard deviations from the mean of latencies (*n* = 6) and trials in which there was no hard cessation of speech (*n* = 80) were removed, leaving 274 trials for consideration. Each trial comprised two electrodes, such that the current flowed into cortex through dipole 1 (“in”) and flowed out of cortex through dipole 2 (“out”). There is no functional difference between stimulation arising from an “in” (i.e., source) or “out” (i.e., sink) electrode. As such, some trials had two electrodes in the region of interest, some trials had electrodes spanning two regions of interest, and some trials had only one electrode within a region of interest. Because our broad regions of interest were sensorimotor cortex (including both precentral and postcentral gyri), IFG, STG, supramarginal gyrus, and MTG, those trials for which stimulation was applied outside these areas (*n* = 23) were removed for statistical analyses, but these trials were preserved in all brain plots. Among these exclusions were two trials from one patient for which a depth electrode spanned precentral gyrus and white cerebral matter. The 251 remaining trials comprised two grid electrodes on cortex with at least one electrode (dipole 1 or dipole 2) in a region of interest. After removal of trials from other regions, those trials in which stimulation induced afterdischarges (*n* = 82) were also removed (see results section). Thus, the final data set for analysis included 169 unique trials in 16 patients. Of the 274 trials analyzed, only nine trials were elicited using the Pledge of Allegiance; following data cleanup, only two of these trials were included in the final statistical analysis. See Table 2 for complete electrode counts in each region from all 274 trials considered for analysis, pooled across trials that induced motor-based arrest and speech arrest. The central column includes electrode counts for trials on which stimulation did not induce afterdischarges (*n* = 185); the right column includes electrode counts for trials on which stimulation induced afterdischarges (*n* = 89). The top section includes electrode counts in the five broad regions of interest, where the top central section includes the electrodes for which stimulation did not induce afterdischarges and that were included in the final analyses (*n* = 165). The histograms in Fig. 1 show the distributions of latencies for all trials (panel B) and for the subset of trials that did not induce afterdischarges (panel C).

**Table 2.** Electrode counts from all 274 trials considered in the present analysis.

| Region | No after-discharges<sup>b</sup> |  | After-discharges |  |
| --- | --- | --- | --- | --- |
|  | in | out | in | out |
| **Regions of interest<sup>a</sup>** |  |  |  |  |
| postcentral | 24 | 24 | 3 | 6 |
| precentral | 35 | 29 | 10 | 9 |
| pars opercularis | 9 | 9 | 12 | 20 |
| pars triangularis | 21 | 16 | 10 | 5 |
| supramarginal | 15 | 47 | 4 | 6 |
| rostral STG | - | - | 8 | 3 |
| middle STG | 17 | 8 | 15 | 14 |
| caudal STG | 29 | 19 | 2 | 6 |
| rostral MTG | - | - | 1 | 1 |
| middle MTG | 4 | 4 | - | - |
| caudal MTG | 11 | 9 | 7 | 5 |
| **Subtotal** | **165** | **165** | 72 | 75 |
| **Other regions** |  |  |  |  |
| caudal MFG | 2 | 3 | 1 | 2 |
| rostral MFG | 4 | 2 | 7 | 3 |
| superior parietal | - | 1 | - | - |
| banks superior temporal sulcus | - | - | 7 | 5 |
| transverse temporal | - | - | 1 | - |
| inferior temporal | 1 | 1 | - | 2 |
| middle temporal | - | - | - | - |
| precentral (depth) | 2 | - | - | - |
| insula | 3 | - | - | - |
| left cerebral white matter | 6 | 8 | - | 1 |
| left pallidum | 1 | - | - | - |
| left putamen | 1 | - | - | - |
| unknown | - | 5 | 1 | 1 |
| **Subtotal** | **20** | **20** | 17 | 14 |
| **Grand total** | **185** | **185** | **89** | **89** |

<sup>a</sup>The top section includes trials within the five broad regions of interest, including trials from dipole 1 (current flowing in) and dipole 2 (current flowing out). Regions include sensorimotor cortex (postcentral/precentral gyrus), IFG (pars opercularis/triangularis), STG (rostral/middle/caudal), supramarginal gyrus, and MTG (rostral/middle/caudal). Columns showing trials with stimulation in other regions are italicized, as these were not included in the present analyses.
<sup>b</sup>The central section includes trials for which stimulation did not induce afterdischarges. Columns showing trials that did induce afterdischarges are italicized, as these were not included in the present analyses.
STG: superior temporal gyrus; MTG: middle temporal gyrus; MFG: middle frontal gyrus

We first used *t*-tests to compare locations and latencies for motor hits versus speech arrest hits, and to compare latencies for trials with versus without afterdischarges. Controlling for stimulation parameters, we then determined whether region (sensorimotor cortex, IFG, STG, supramarginal gyrus, MTG) predicts adjusted latency from stimulation onset to speech arrest using a non-parametric Kruskal-Wallis test. To determine whether latencies were different between the five broad regions, we conducted post-hoc two-sided rank sum Wilcoxon tests between each pair of regions while controlling for multiple comparisons. Statistical tests were performed using functions in the base “stats” packages in R, including ‘t.test,’ ‘kruskal.test,’ and ‘wilcox.test.’ Finally, latency patterns were visualized on brain regions across cortex using custom Matlab^26^ code, and then compared using smoothed density plots via the “ggplot2” package in R. Bootstrapped medians and confidence intervals from 5000 resampled iterations of our data were computed using the ‘groupwiseMedian’ function in the “rcompanion” package in R.

### Data availability

Anonymized data will be shared upon request.

## Results

### Motor versus speech arrest hits in five broad regions

In order to understand the spatial topography of speech arrest, we first analyzed the spatial distribution of motor hits versus speech arrest hits and verified locations (Fig. 2). We found that the main regions implicated included our broad regions of interest: sensorimotor cortex, IFG, STG, and supramarginal gyrus, in addition to MTG. Regardless of type of arrest elicited, latencies (i.e., the “adjusted latencies” described in the methods section) for motor hits were not different from latencies for speech arrest hits across all trials including afterdischarges, *t*(249) = 0.770, *p* = 0.442 (Fig. 2A). Upon removal of all trials with afterdischarges, we still found no difference between motor-based arrest versus speech arrest hits despite there being fewer trials, *t*(167) = 1.48, *p* = 0.141 (Fig. 2B).

**Figure 2.**
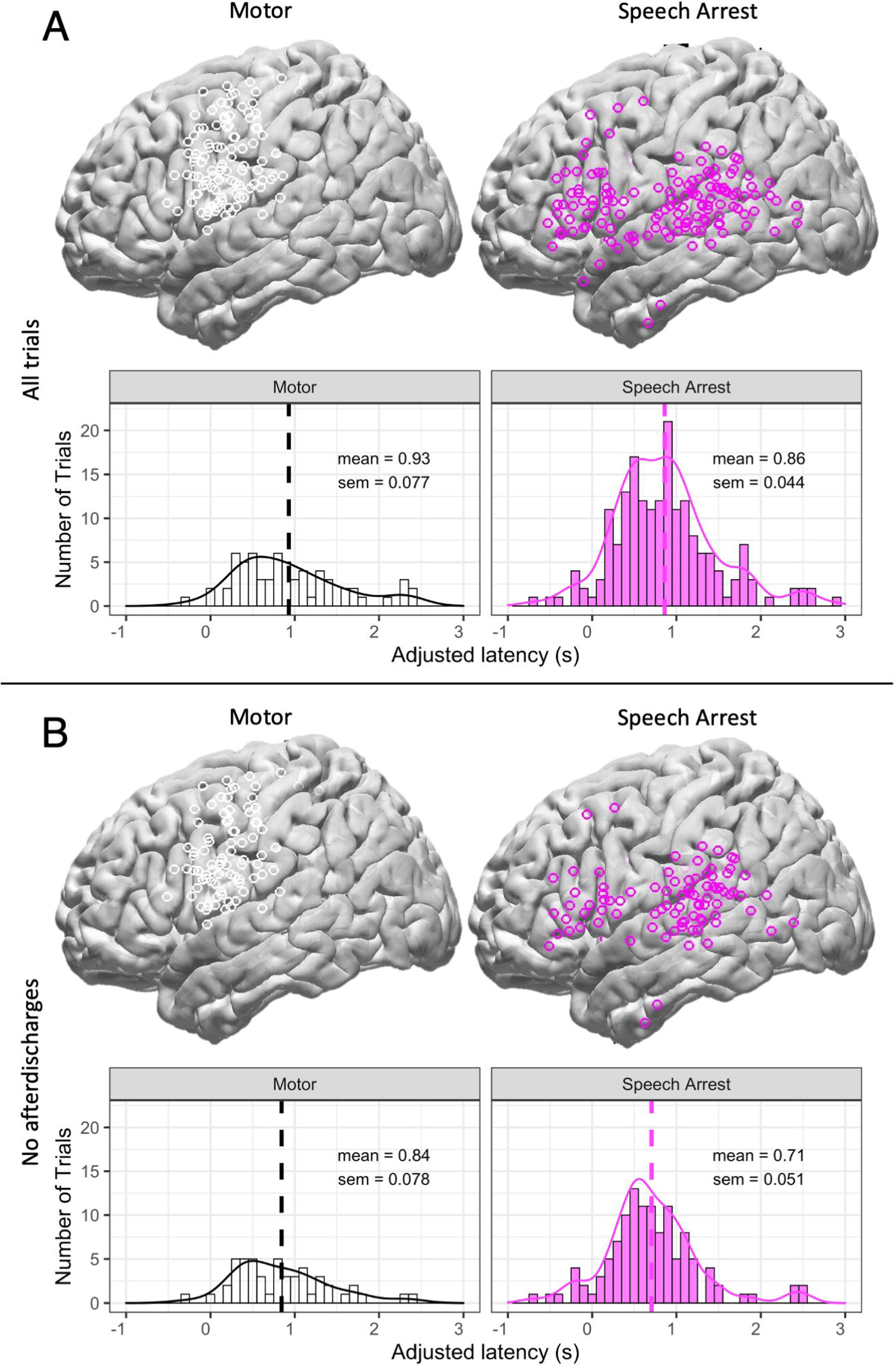
Cortical locations and distributions of latencies for all motor and speech arrest hits, controlling for trials with afterdischarges. **(A)** Brain plots of cortical locations (top) and histograms and density plots of latencies (bottom) for all motor hits (left) and speech arrest hits (right) in all trials including those with and without afterdischarges. The dashed lines in the histograms indicate the mean, which is labeled with the standard error of the mean. A two-sample t-test indicated that latencies for motor hits (*n* = 65) were not different from latencies for speech arrest hits (*n* = 186) across all trials including afterdischarges, *t*(249) = 0.770, *p* = 0.442. **(B)** Same as panel A, but only those trials excluding afterdischarges are shown. A two-sample t-test indicated that latencies for motor hits (*n* = 50) were not different from latencies for speech arrest hits (*n* = 119) across all trials excluding afterdischarges, *t*(167) = 1.48, *p* = 0.141.

### Comparing trials with and without afterdischarges

In order to rule out the effect of epileptic activity, we removed all trials with afterdischarges from subsequent analyses. Comparison of latencies for all motor hits and speech arrest hits in trials with and without afterdischarges provides further rationale for this exclusion (Fig. 3). That is, trials with afterdischarges had longer latencies than trials without afterdischarges for motor hits, *t*(63) = -2.06, *p* = 0.0437 (Fig. 3A) and for speech arrest hits, *t*(184) = -4.97, *p* < 0.0001 (Fig. 3B).

**Figure 3:**
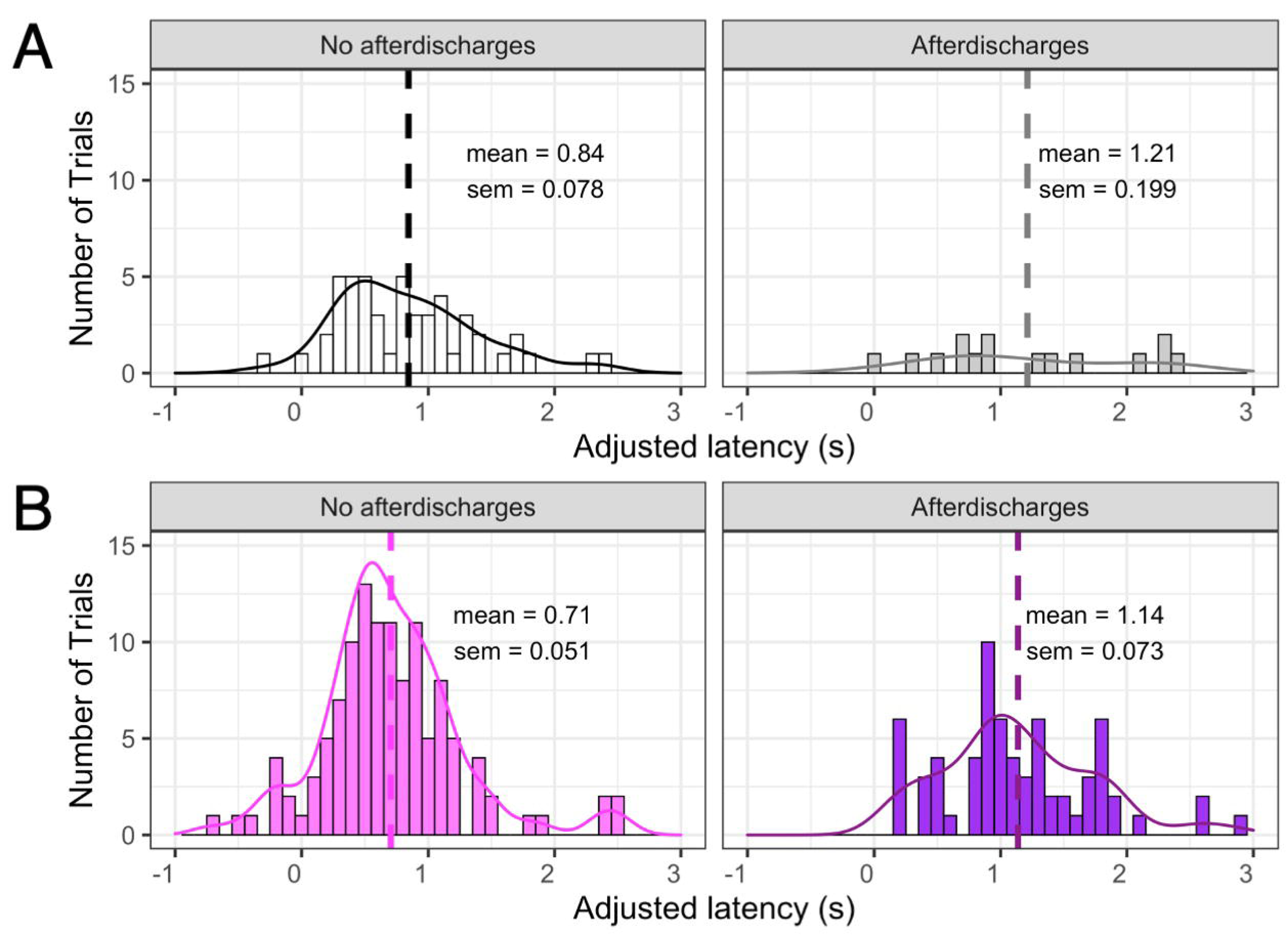
Distributions of latencies for all motor hits, controlling for trials with afterdischarges. **(A)** Histograms and density plots of latencies for all motor hits, including trials with no afterdischarges (left) and trials with afterdischarges (right). Dashed lines indicate the mean, which is labeled with the standard error of the mean. A two-sample t-test indicated that latencies for trials with afterdischarges (*n* = 15) had longer latencies than trials without afterdischarges (*n* = 50) for motor hits, *t*(63) = -2.06, *p* = 0.0437. **(B)** Same as panel A, but only speech arrest hits are shown. A two-sample t-test indicated that latencies for trials with afterdischarges (*n* = 67) had longer latencies than trials without afterdischarges (*n* = 119) for speech arrest hits, *t*(184) = -4.97, *p* < 0.0001.

### Regional differences in latency

Before comparing regional differences in latency, we considered distributions of motor hits and speech arrest hits in each cortical region (Fig. 4A). The majority of motor hits were in sensorimotor cortex (87.5% of 96 instances), where all but 12 instances were in precentral gyrus (53.1%) or postcentral gyrus (34.4%). In contrast, speech arrest hits were relatively more spread out across cortex. Within sensorimotor cortex, there were relatively more motor hits (75%) than speech arrest hits (25%); both motor hits and speech arrest hits tended to have roughly equal portions of electrodes from precentral gyrus and postcentral gyrus. Within sensorimotor cortex, motor hits tended to span both dorsal and ventral regions, whereas speech arrest hits tended to be isolated to ventral regions (see discussion). In other regions, there were substantially more speech arrest hits relative to the number of motor hits within supramarginal gyrus (96.8%), STG (94.5%); MTG (100%), and IFG (89%).

**Figure 4:**
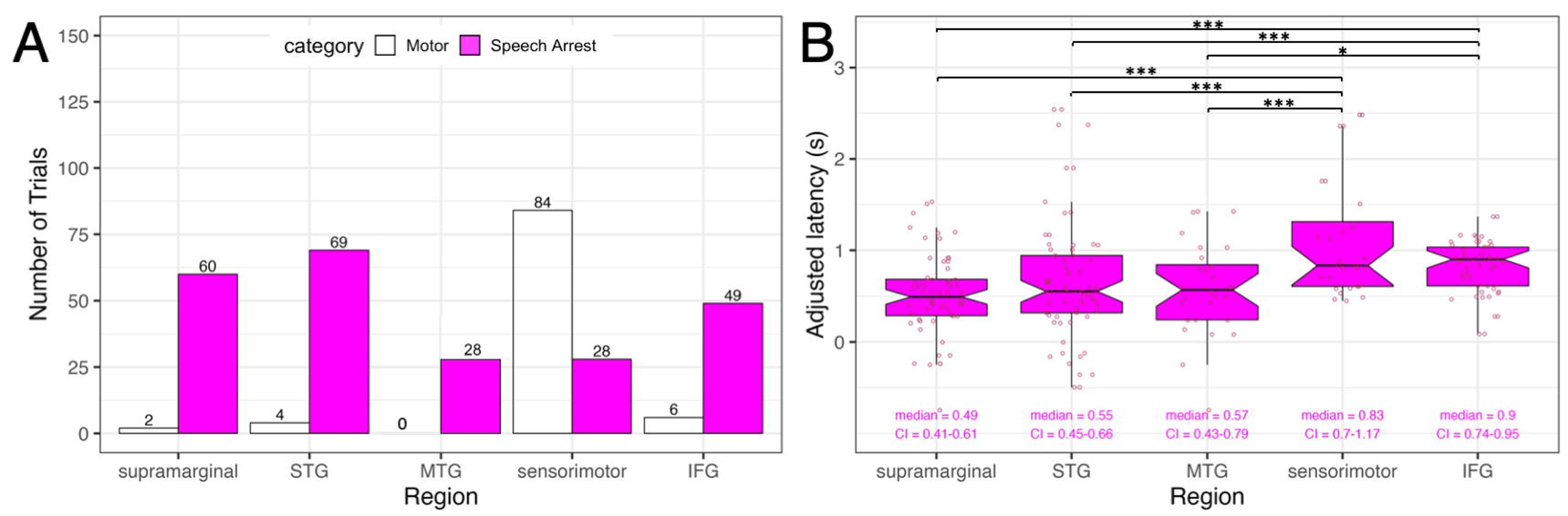
Summary of motor hits and speech arrest hits in each cortical region. **(A)** Counts of motor hits and speech arrest hits in each cortical region, with the number of electrodes in each group labeled. **(B)** Boxplots of latencies for each cortical region, with individual data points shown and labels indicating median latency with 95% confidence intervals for the median labeled and shown as notches. Three asterisks indicate significant post-hoc tests at the corrected *p* = 0.0109 level; single asterisks indicate marginal significance at the 0.05 level. STG: superior temporal gyrus; MTG: middle temporal gyrus; IFG: inferior frontal gyrus

Including all trials of speech arrest, we tested statistically whether latencies in each broad cortical region differed from one another (Fig. 4B). A nonparametric Kruskal-Wallis test indicated that variances between the five broad cortical regions were significantly different from one another, x^2^(4) = 28.798, *p* < 0.00001. To determine whether stimulation parameters influenced latency differently for each region, we ran a linear mixed effects regression model predicting latency on a subset of the data including only the final trial in each stimulation sequence, so that the highest electrical stimulation level correctly corresponded with the final trial in that sequence. We predicted latency from region, stimulation level (in mA), and the interaction between these two predictors, while including a random intercept for patient. Region was a significant predictor of latency [F(4, 81.872) = 2.759, p = 0.0331]; neither stimulation level [F(4, 81.188) = 1.359, p = 0.247] nor the interaction between region and stimulation level [F(4, 81.609) = 1.171, p = 0.330] significantly predicted latency. We conducted ten post-hoc two-sided rank sum Wilcoxon tests for which we corrected for multiple comparisons using False Discovery Rate^30^ (q = 0.05) providing a corrected threshold p-value of 0.0109. Latencies in supramarginal gyrus were significantly shorter than in Sensorimotor Cortex (*W* = 357, *p* = 0.0000157) and in IFG (*W* = 796, *p* = 0.00004081). Latencies in STG were significantly shorter than in Sensorimotor Cortex (*W* = 560, *p* = 0.00124) and in IFG (*W* = 1186, *p* = 0.00591), and latencies in MTG were significantly shorter than in Sensorimotor Cortex (*W* = 204, *p* = 0.00218). Latencies in MTG were shorter than in IFG (*W* = 445, *p* = 0.0109), but these differences were significant only at the uncorrected p-value threshold of 0.05. Latencies between all other regions were not significantly different from one another (Sensorimotor Cortex versus IFG [*W* = 593.5, *p* = 0.3298]; supramarginal gyrus versus STG [*W* = 1833.5, *p* = 0.265]; supramarginal gyrus versus MTG [*W* = 764, *p* = 0.499], STG versus MTG [*W* = 996, *p* = 0.814].

### Temporal map of latencies across cortex

Finally, we analyzed the temporal dynamics of latencies to speech arrest across cortex in four time bins (Fig. 5A). Latencies less than 0.5 seconds were associated with a high density of speech arrest following stimulation in supramarginal gyrus and STG. At latencies from 0.5-1 second, the density of speech arrest in STG faded while the density in IFG increased. For latencies greater than 1 second, there was relatively high density of speech arrest in IFG and possibly also in STG.

**Figure 5:**
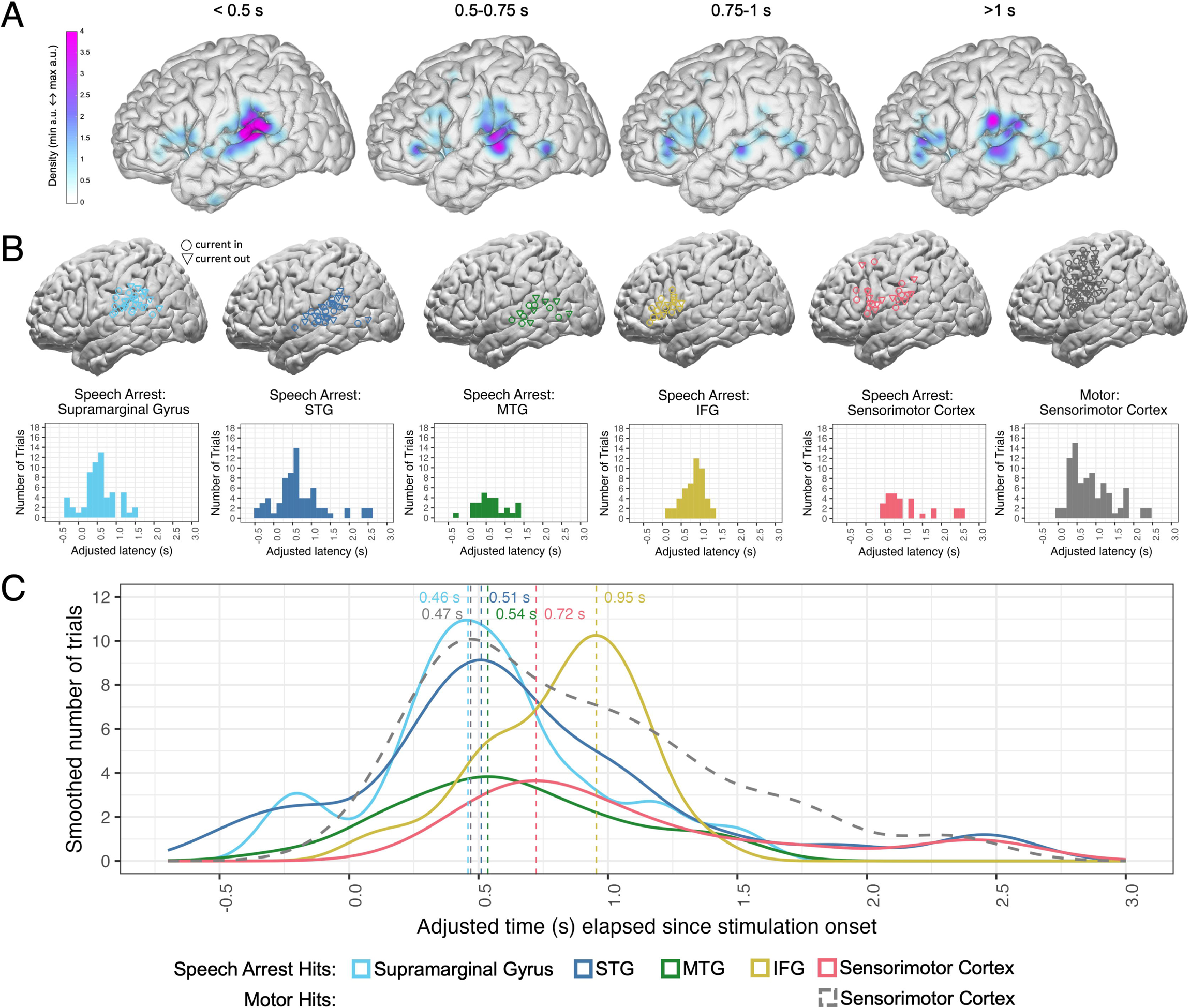
Cortical locations and distributions of speech arrest hits relative to motor hits. **(A)** Map of the density of speech arrest hits across cortex in four time bins of adjusted latency. **(B)** Cortical locations and distributions of adjusted latencies for all speech arrest hits within each region, and for all motor hits in sensorimotor cortex. All electrodes are based on within-subject anatomy, which may differ in plotting location following normalization to Montreal Neurological Institute (MNI) coordinates. Circles indicate electrodes through which the current was flowing in (dipole 1) while the triangles indicate electrodes through which the current was flowing out (dipole 2). **(C)** Smoothed distributions of adjusted latencies, including speech arrest hits in supramarginal gyrus (light blue), STG (dark blue), MTG (green), IFG (yellow) and sensorimotor cortex (coral), and as a comparison, the motor hits in sensorimotor cortex (dark gray dashed distribution). Density plots are scaled based on the number of electrodes in each distribution and a 0.15 second bin width. Dashed vertical lines mark the peak of each smoothed distribution line, with labeled peak values. STG: superior temporal gyrus; MTG: middle temporal gyrus; IFG: inferior frontal gyrus

We also analyzed the temporal dynamics of speech arrest across cortex by looking at distributions of latencies in the broad regions of interest, with the cortical locations and distribution of latencies of motor hits in sensorimotor cortex also plotted for reference (Fig. 5B). Looking at speech arrest patterns across the regions (Fig. 5C), we report here the peak latency from the distribution of latencies for each region, accompanied by the bootstrapped median (5000 iterations) and 95% confidence intervals for the median. There are early peaks in temporoparietal areas, including supramarginal gyrus at 0.46 seconds (median = 0.49, CI = 0.41-0.61), STG at 0.51 seconds (median = 0.55, CI = 0.45-0.66), and MTG at 0.54 seconds (median = 0.57, CI = 0.43-0.79). Later peaks occur in sensorimotor cortex at 0.72 seconds (median = 0.83, CI = 0.70-1.17) and in IFG at 0.95 seconds (median = 0.90, CI = 0.74-0.95). In contrast to the later peaks for speech arrest in sensorimotor cortex, motor hits in sensorimotor cortex show an early peak at 0.47 seconds (median = 0.79, CI = 0.55-0.92). Additionally, Fig. 5B shows how motor hits span sensorimotor cortex whereas speech arrest hits in this region tend to be ventrally located. See Supplemental Figs. S2 and S3 depicting an alternate analysis in which sensorimotor cortex is divided anatomically into precentral and postcentral gyri.

## Discussion

By reporting on the timing of stimulation to speech arrest involving the spatial precision of electrocorticography, we observed that shorter latencies were associated with motor-based speech interruptions across sensorimotor cortex and with speech arrest in supramarginal gyrus and STG. In contrast, longer latencies were associated with speech arrest in ventral portions of precentral and postcentral gyri in sensorimotor cortex and in IFG. These results suggest that stimulation in specific neural regions interrupts disparate processes of speech motor control with execution and planning occurring in distinct areas within sensorimotor cortex.

The goal of this study was to characterize patterns of region-specific latency from stimulation onset to motor-based speech interruptions and to speech arrest in five peri-sylvian cortical regions. Trials with afterdischarges were excluded from analyses because epileptic activity could not be ruled out in these trials and because speech arrest latencies that induced afterdischarges were significantly longer than trials that did not induce afterdischarges (Fig. 3). We believe that the afterdischarges contributed to an interruption in the speech network beyond the region-specific interruptions induced by DES, likely interacting with epileptiform activity.

In summary, there were relatively more motor hits in sensorimotor regions than in other cortical areas, while there were relatively more speech arrest hits in the other cortical areas, including supramarginal gyrus, STG, MTG, and IFG (see Fig. 4A). Latencies for electrodes that were labeled as motor hits were not different than latencies for speech arrest hits (Fig. 2). For speech arrest, the earliest latencies were in supramarginal gyrus and STG, and latencies in these regions (and in MTG to a lesser extent) were significantly shorter than latencies in both sensorimotor cortex and IFG. Latencies for speech arrest within supramarginal gyrus, STG, and MTG were not different from one another; latencies in sensorimotor cortex and IFG were also not different from one another, contrary to our prediction.

We had hypothesized that stimulation in sensorimotor cortex would interrupt motor execution midstream leading to the prediction that stimulation within this region would cause the shortest latencies. We were surprised to find that the earliest peak latencies for speech arrest were in supramarginal gyrus (0.46 s) and STG (0.51 s), and that latencies in these regions were significantly shorter than latencies in sensorimotor cortex (peak at 0.72 s) and in IFG (peak at 0.95 s). The early peak latencies for speech arrest in supramarginal gyrus and STG aligned with the early peak latency for motor hits in sensorimotor cortex (0.47 s), suggesting that temporoparietal and sensorimotor areas may be involved with interruptions in the motor execution phase of speech production.

In contrast, we had hypothesized that IFG would interrupt motor planning leading to the prediction that stimulation in this region would cause relatively longer latencies to speech arrest. The relatively long latencies that we observed for speech arrest in both sensorimotor cortex and in IFG suggest that there was interruption to planning processes in both cortical areas. Within sensorimotor cortex, latencies to motor-based speech arrest occurred relatively early and within areas throughout ventral and dorsal portions of precentral and postcentral gyri. However, latencies to events labeled as speech arrest occurred relatively late and within IFG and more ventrally located parts of sensorimotor cortex. This novel finding suggests that motor execution and planning processes occur in distinct areas within sensorimotor cortex. However, we acknowledge that latencies we measured in postcentral gyrus were significantly longer than those in precentral gyrus (see Supplemental Figs. S2 and S3, which we discuss further below).

What stands out in this study is the surprisingly short latencies to speech arrest found in supramarginal gyrus and STG. We considered two potential mechanisms for these quick interruptions in temporoparietal areas. First, stimulation could interfere with lexical access when speaking, as previous research has shown that both speech arrest and anomia occur within supramarginal and STG sites.^6^ However, when reviewing the sites for which anomia was elicited in our data set, this hypothesis was not supported because the hits for speech arrest in supramarginal gyrus and STG were not also hits for anomia. Instead, our data support a self-monitoring hypothesis in which stimulation causes an interruption within the auditory feedback loop, thus interrupting real-time feedback-based updates to the outgoing motor plan. This claim is supported by our finding that latencies for speech arrest in STG were comparable to latencies to motor-based speech interruptions in sensorimotor cortex. This claim is also supported by research linking deficits in auditory-verbal short-term memory to supramarginal gyrus.^31^ Furthermore, multiple studies have shown efference copy in temporal regions,^32,33^ a process by which motor regions inform auditory areas of planned outgoing motor commands. Thus, this link between motor and auditory areas might explain the similar latencies observed between sensorimotor cortex and STG.

Our data corroborate past findings from multiple studies reporting speech arrest in two main cortical regions, including sensorimotor cortex (both precentral and postcentral gyri) and IFG.^1,6-8^ As the most consistently reported site for speech arrest is precentral gyrus, our study highlights how most of the speech interruptions in this region were not speech arrest per se because they were determined to be motoric in nature. Our findings also reveal robust speech arrest in IFG, in contrast to studies showing sparse speech arrest in this region,^5,9^ but compatible with other reports.^6-8,34^ We also showed robust speech arrest in temporoparietal areas, in contrast to past reports indicating sparse and highly variable speech arrest in supramarginal gyrus^6^ and in STG.^5-7,9^ As reported in Table 2, most speech interruptions occurred in our five regions of interest across the sixteen patients included in the analysis, suggesting that we have provided a comprehensive analysis of speech arrest across cortex.

We now acknowledge the limitations to the present study. The main limitation concerns electrode coverage. Coverage was dictated by clinical necessity and although our data set collectively included extensive coverage of sensorimotor cortex, IFG, supramarginal gyrus, STG, and MTG, some patients did not have coverage of all five regions of interest. Secondly, our functional definition of sensorimotor cortex included electrodes from both precentral and postcentral anatomical regions, as supported in other research.^13-15^ However, we report an alternate analysis in Supplemental Figs. S2 & S3 in which it becomes apparent that latencies to speech arrest in postcentral gyrus (*n* = 15) were longer than those in precentral gyrus (*n* = 13). Even though longer latencies in postcentral gyrus may contribute disproportionately to our claim that sensorimotor regions have longer latencies, electrodes in precentral gyrus also tended to have longer latencies than temporoparietal regions (supramarginal gyrus, in particular). Hence, we believe that a larger data set might reveal precentral gyrus (like its functional counterpart, postcentral gyrus) as having significantly longer latencies to speech arrest. A third limitation regards our inability to make claims about lateralization, as the vast majority of patients undergo language mapping only on the left hemisphere. Only one patient who was initially considered had coverage over the right hemisphere; this coverage included seven trials that had afterdischarges, so after data cleanup, this patient was no longer included the final analysis. Therefore, our lack of coverage of the right hemisphere limits our capacity to answer any questions related to lateralization. A fourth limitation is that we could not control for stimulation level for all trials in our data set because the documented stimulation level corresponded only to the final trial in each stimulation sequence. However, for the subset of our data that included only these final trials, stimulation level did not predict latency, nor did it correspond with latencies in a region-specific manner. Although it is unlikely that stimulation level affected latency differently within distinct cortical regions, future research should test this directly by documenting stimulation level for each trial within a stimulation sequence.

Most of the previous studies reporting cortical sites for inducing speech arrest were intra-operative and therefore had limited cortical coverage.^1,6-9^ Our study complements a growing number of reports examining speech arrest mapped extra-operatively.^5,35^ We believe that the broader cortical coverage enabled by extra-operative DES mapping lends strong support to the robustness of the spatial patterns observed in our data set. Furthermore, electrode diameters and inter-electrode distances used during each approach differ in an important way. Cortical grids used in extra-operative mapping contain bipolar electrodes that are 2.4 millimeters in diameter with ten-millimeter inter-electrode distances, whereas cortical probes used intra-operatively contain electrodes that are one millimeter in diameter with five-millimeter inter-electrode distances.^36^ These differences in DES mapping approach suggest that even though extra-operative mapping has broader coverage across cortex, intra-operative mapping could have more detailed spatial sampling within the probed cortical regions. This suggests that arriving at a consensus of insights derived through both approaches will be important to understanding the spatial and temporal dynamics of speech arrest across cortex.

This study provides a unique window into the functional architecture and neural timing of speech arrest by providing a map of how long it takes stimulation to induce speech arrest across cortical regions. Our study corroborates past reports of robust speech arrest in precentral gyrus and IFG while lending strong support to supramarginal gyrus, STG, MTG, and postcentral gyrus also being robust sites for speech arrest. Analyses of timing from stimulation onset to speech arrest revealed that supramarginal gyrus and STG had the shortest latencies, comparable to those from motor-based arrest induced broadly in sensorimotor cortex. In contrast, speech arrest induced in ventral parts of sensorimotor cortex (both precentral and postcentral gyri) and in IFG had relatively long latencies. These results are compatible with a speech production framework in which longer latencies are associated with speech motor planning (e.g., IFG), whereas shorter latencies are associated with motor execution (e.g., sensorimotor cortex), which encompasses real-time feedback-based updates to motor execution. Clinically, our results suggest that stimulation in sensorimotor cortex generally yields shorter latencies suggesting an interruption to motor execution, but that stimulation in ventral sensorimotor regions as well as in IFG yields relatively long latencies suggesting an interruption to planning processes.

## Acknowledgements

The authors acknowledge the Finding a Cure for Epilepsy and Seizures (FACES) Foundation and our patients for participating in research.

## Funding

This research was supported by the National Institute on Deafness and Other Communication Disorders Grants F32DC021094 (H. Kabakoff, PI) and R01DC018805 (A. Flinker, PI), and the National Institute of Neurological Disorders and Stroke Grants R01NS109367 (A. Flinker, PI) and R01NS115929 (A. Flinker, PI).

## Competing interests

Author Dr. Daniel Friedman receives salary support for consulting and clinical trial related activities performed on behalf of The Epilepsy Study Consortium, a non-profit organization. Dr. Friedman receives no personal income for these activities. NYU receives a fixed amount from the Epilepsy Study Consortium towards Dr. Friedman’s salary. Within the past two years, The Epilepsy Study Consortium received payments for research services from: Biohaven, BioXcell, Cerevel, Cerebral, Epilex, Equilibre, Jannsen, Lundbeck, Praxis, Puretech, Neurocrine, SK Life Science, Supernus, UCB, and Xenon. Dr. Friedman has also served as a paid consultant for Neurelis Pharmaceuticals, holds equity interests in Neuroview Technology, and received royalty income from Oxford University Press. All other authors report no competing interests.

## Supplemental material

Supplementary Figs. S1, S2, and S3 are available at *Brain Communications* online.

## Notes

### Summary of Updates

We have made the edits as suggested by peer reviewers, including the clarification of our hypotheses, an adjustment to the statistical analyses used, the addition of a new online supplement with an alternate analysis, and the addition of text revision as needed throughout.

