## Supplemental material for "Timing and location of speech errors induced by direct cortical stimulation"

**A**

**B**

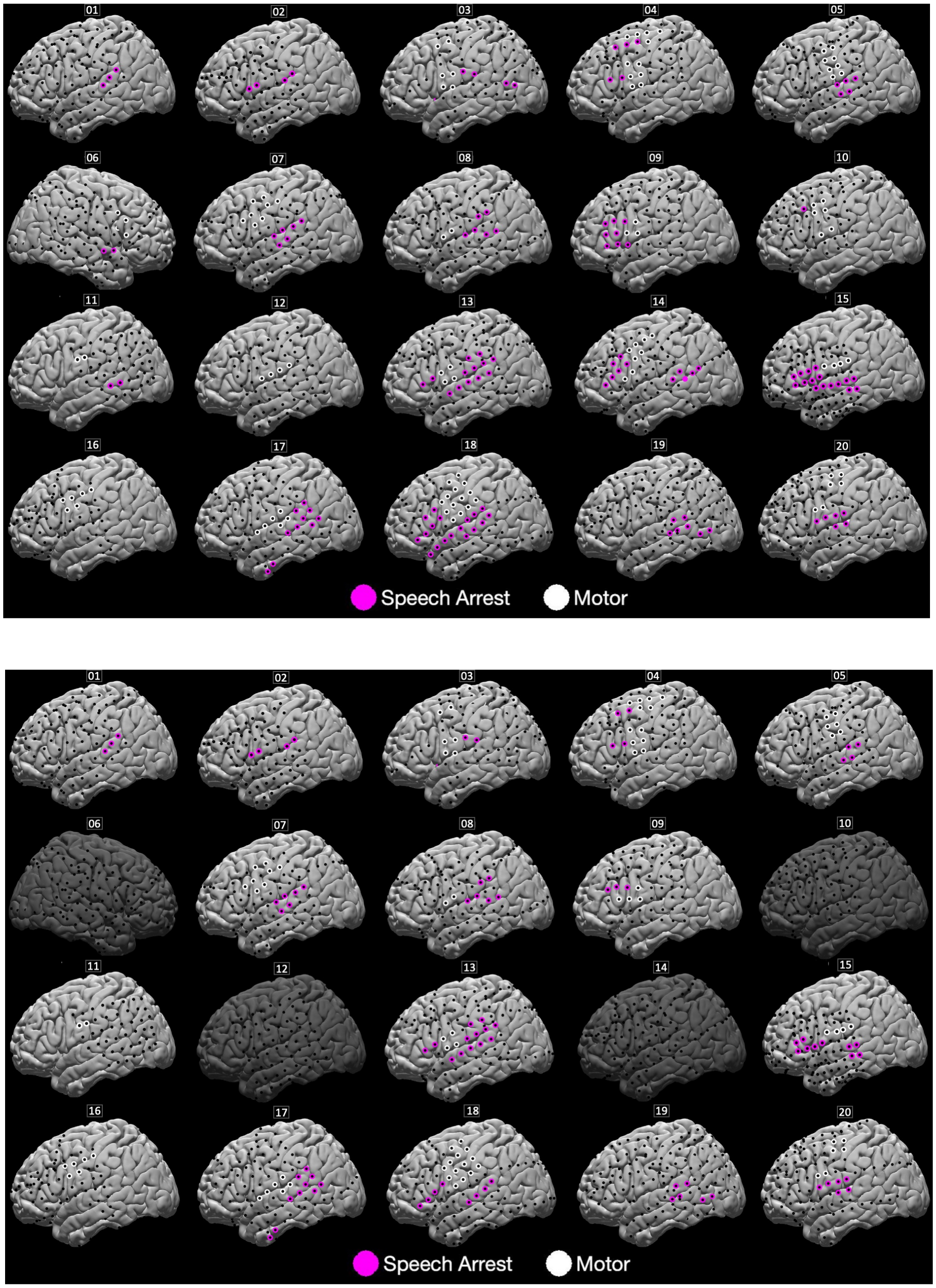


**Figure S1: Individual patient motor-based hits and speech arrest hits. (A)** Electrode locations across cortex for patients 1-20, including trials with afterdischarges. **(B)** Electrode locations across cortex for 16 patients following removal of trials with afterdischarges. For both panels, white highlighting around electrode location indicates that stimulation induced motor-based arrest at that site; magenta highlighting around electrode location indicates that stimulation induced speech arrest at that site. All electrode locations are based on within-subject anatomy, which may differ in plotting location following normalization to Montreal Neurological Institute (MNI) coordinates.

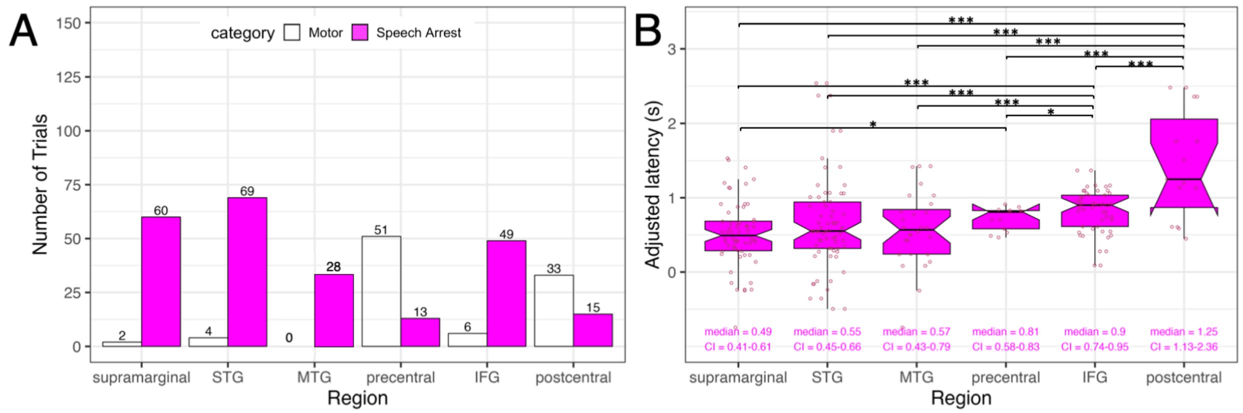


**Figure S2: Summary of motor hits and speech arrest hits in each cortical region, with precentral gyrus and postcentral gyrus listed as separate regions. (A)** Counts of motor hits and speech arrest hits in each cortical region, with the number of electrodes in each group labeled. **(B)** Boxplots of latencies for each cortical region, with individual data points shown and labels indicating median latency with 95% confidence intervals for the median labeled and shown as notches. Three asterisks indicate significant post-hoc tests at the corrected $p = 0.0240$ level; single asterisks indicate marginal significance at the 0.05 level.
STG: superior temporal gyrus; MTG: middle temporal gyrus; IFG: inferior frontal gyrus

Including all trials of speech arrest, we tested statistically whether latencies in each broad cortical region differed (Fig. S2B). A nonparametric Kruskal-Wallis test indicated that variances between the six broad cortical regions were significantly different from one another, $\chi^2(5) = 33.906$, $p < 0.00001$. We conducted fifteen post-hoc two-sided rank sum Wilcoxon tests for which we corrected for multiple comparisons using False Discovery Rate[28] (q = 0.05) providing a corrected threshold p-value of 0.0240.

- Latencies in supramarginal gyrus were significantly shorter than in postcentral gyrus ($W = 124.5$, $p = 0.0000167$) and in IFG ($W = 796$, $p = 0.0000408$). Latencies in supramarginal gyrus were shorter than in precentral gyrus [$W = 233$, $p = 0.0240$], but this difference was significant only at the uncorrected p-value threshold of 0.05.

- Latencies in STG were significantly shorter than in postcentral gyrus ($W = 208$, $p = 0.000307$) and in IFG ($W = 1186$, $p = 0.00591$).
- Latencies in MTG were significantly shorter than in postcentral gyrus ($W = 68$, $p = 0.000310$) and in IFG ($W = 445$, $p = 0.0109$).
- Latencies in precentral gyrus were significantly shorter than in postcentral gyrus ($W = 40$, $p = 0.00861$). Latencies in precentral gyrus were shorter than in IFG [$W = 421.5$, $p = 0.0762$], but this difference was significant only at the uncorrected p-value threshold of 0.05.
- Latencies in IFG were significantly shorter than in postcentral gyrus ($W = 172$, $p = 0.00199$).
- Latencies between all other regions were not significantly different from one another (supramarginal gyrus versus STG [$W = 1833.5$, $p = 0.265$]; supramarginal gyrus versus MTG [$W = 764$, $p = 0.499$], STG versus MTG [$W = 996$, $p = 0.814$], STG versus precentral gyrus [$W = 352$, $p = 0.229$], and MTG versus precentral gyrus [$W = 136$, $p = 0.2022$].

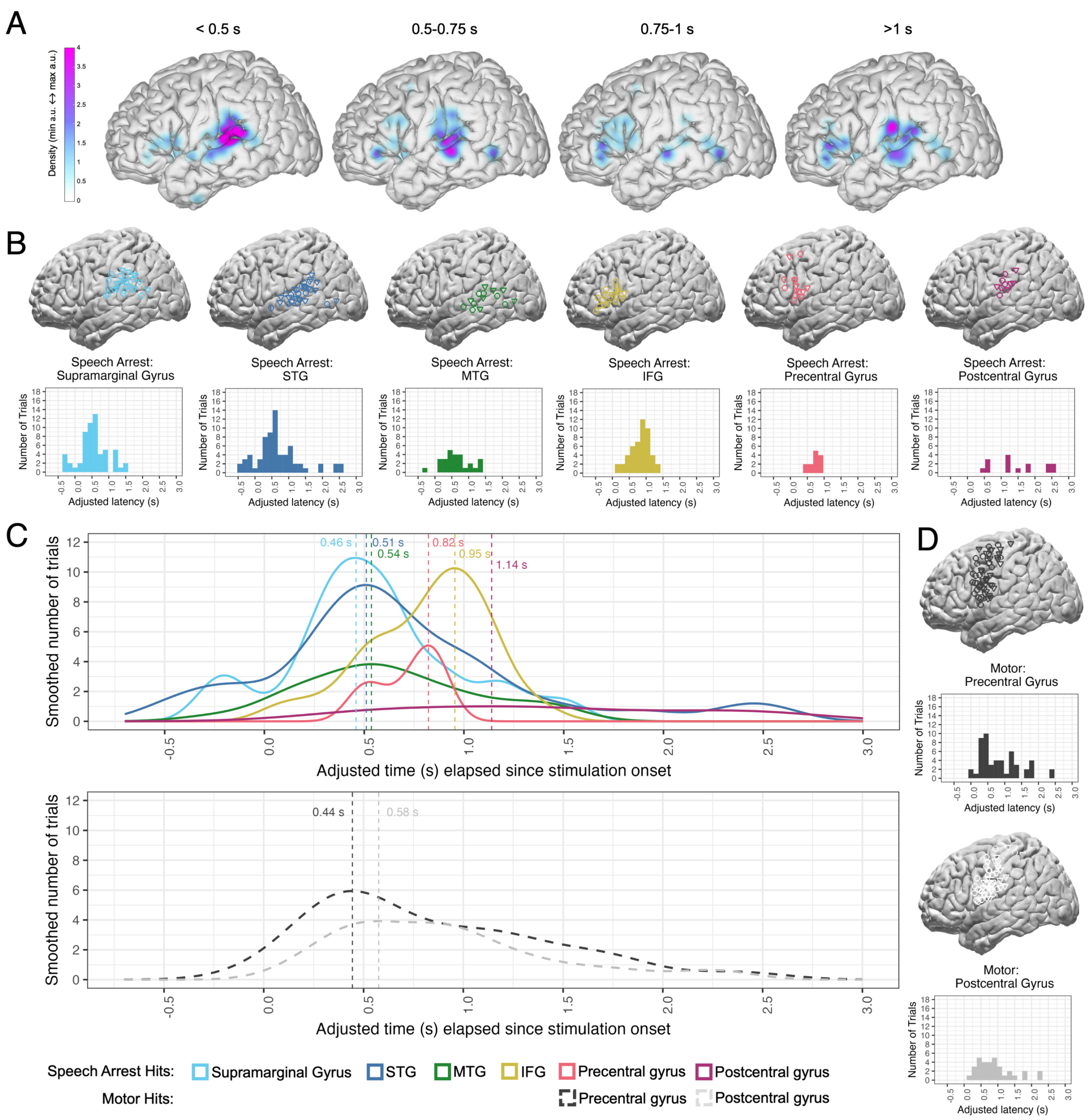


**Figure S3*: Cortical locations and distributions of speech arrest hits relative to motor hits, with precentral gyrus and postcentral gyrus listed as separate regions.* (A)** Map of the density of speech arrest hits across cortex in four time bins of adjusted latency. **(B)** Cortical locations and distributions of adjusted latencies for all speech arrest hits within each region, and for all motor hits in precentral and postcentral gyri. All electrodes are based on within-subject anatomy, which may differ in plotting location following normalization to Montreal Neurological Institute (MNI) coordinates. **(C)** Smoothed distributions of adjusted latencies, including speech arrest hits in supramarginal gyrus (light blue), STG (dark blue), MTG (green), IFG (yellow), precentral gyrus (coral), and postcentral gyrus (fuchsia), and as a comparison, the motor hits in precentral gyrus (dark gray dashed distribution) and postcentral gyrus (light gray dashed distribution). Density plots are scaled based on the number of electrodes in each distribution and a 0.15 second bin width. Dashed vertical lines mark the peak of each smoothed distribution line, with labeled peak values. STG: superior temporal gyrus; MTG: middle temporal gyrus; IFG: inferior frontal gyrus

Looking at speech arrest patterns across the regions (Fig. S3C), we report here the peak latency from the distribution of latencies for each region, accompanied by the bootstrapped median (5000 iterations) and 95% confidence intervals for the median. There are early peaks in temporoparietal areas, including supramarginal gyrus at 0.46 seconds (median = 0.49, CI = 0.41-0.61), STG at 0.51 seconds (median = 0.55, CI = 0.45-0.66), and MTG at 0.54 seconds (median = 0.57, CI = 0.43-0.79). Later peaks occur in precentral gyrus at 0.82 seconds (median = 0.81, CI = 0.58-0.83), IFG at 0.95 seconds (median = 0.90, CI = 0.74-0.95), and postcentral gyrus at 1.14 seconds (median = 1.25, CI = 1.13-2.36). In contrast to the later peaks for speech arrest in precentral and postcentral gyri, motor hits in precentral and postcentral gyri show early peaks at 0.44 seconds (median = 0.70, CI = 0.49-0.97) and 0.58 seconds (median = 0.81, CI = 0.55-0.96), respectively.
